# Metabolic reprogramming from glycolysis to amino acid utilization in cardiac HIF1α deficient mice

**DOI:** 10.1101/579433

**Authors:** Ivan Menendez-Montes, Beatriz Escobar, Beatriz Palacios, Manuel J. Gomez, Elena Bonzon, Alessia Ferrarini, Ana Vanessa Alonso, Luis Jesus Jimenez-Borreguero, Jesus Vázquez, Silvia Martin-Puig

## Abstract

**Rationale:** Hypoxia is an important environmental cue implicated in several physiopathological processes, including heart development. Several mouse models of activation or inhibition of hypoxia have been previously described. While gain of function models have been extensively characterized and indicate that HIF1 signaling needs to be tightly regulated to ensure a proper cardiac development, there is lack of consensus in the field about the functional outcomes of HIF1α loss.

**Objective:** In this study, we aim to assess the consequences of cardiac deletion of HIF1α during heart development and identify the cardiac adaptations to HIF1 loss.

**Methods and Results:** Here, we used a conditional deletion model of *Hif1a* in NKX2.5^+^ cardiac progenitors. By a combination of histology, electron microscopy, massive gene expression studies, proteomics, metabolomics and cardiac imaging, we found that HIF1α is dispensable for cardiac development. *Hif1a* loss results in glycolytic inhibition in the embryonic heart without affecting normal cardiac growth. However, together with a premature increase in mitochondrial number by E12.5, we found global upregulation of amino acid transport and catabolic processes. Interestingly, this amino acid catabolism activation is transient and does not preclude the normal cardiac metabolic switch towards fatty acid oxidation (FAO) after E14.5. Moreover, *Hif1a* loss is accompanied by an increase in ATF4, described as an important regulator of several amino acid transporters.

**Conclusions:** Our data indicate that HIF1α is not required for normal cardiac development and suggest that additional mechanisms can compensate *Hif1a* loss. Moreover, our results reveal the metabolic flexibility of the embryonic heart at early stages of development, showing the capacity of the myocardium to adapt its energy source to satisfy the energetic and building blocks demands to achieve normal cardiac growth and function. This metabolic reprograming might be relevant in the setting of adult cardiac failure.

## INTRODUCTION

The heart is the first organ to form during development, as it is essential to deliver oxygen and nutrients to embryonic tissues from early stages. Different subsets of cardiac progenitors proliferate, migrate and differentiate into the diverse cell types that form the mature heart [1, 2]. Despite of the existence of several types of cardiovascular precursors, NKX2.5 progenitors give rise to the majority of cardiac cells, contributing to the three main heart layers: epicardium, myocardium and endocardium [3]. However, cardiogenesis can result in malformations, with congenital heart defects being present in 1% of live births. Several factors have been involved in developmental cardiac failure, and among them hypoxia, or low oxygen tensions, has been described as an environmental factor of cardiac malformations during pregnancy [4]. Hypoxia-Inducible Factors (HIFs) are known to mediate a well-characterized transcriptional response to hypoxia. HIFs heterodimers are formed by a constitutively expressed β subunit (HIFβ or ARNT) and an oxygen-regulated α subunit, with three different isoforms (1α, 2α and 3α) [5]. Under normoxic conditions, the oxygen sensors Prolyl Hydroxylases (PHDs) hydroxylate HIFα in specific proline residues [6]. This modification is a recognition mark for the von Hippel Lindau (VHL)/E3 ubiquitin ligase complex, that polyubiquitinates and drives α subunits to proteasomal degradation. In hypoxic conditions, α subunits evade degradation due to the inhibited PHD activity, dimerize with β subunits and mediate the adaptive response to hypoxia by activating the transcriptions of their targets genes [7].

In addition to the existence of hypoxic areas during heart development [4, 8], it has been described that chronic exposure of pregnant females to 8% oxygen causes embryonic myocardial thinning, epicardial detachment and death [9] and global deletion of different components of the hypoxia pathway display cardiovascular phenotypes [10]. Moreover, genetic-based overactivation of HIF signaling by inactivation of *Vhl* gene in cardiac cells using different drivers (Mlc2vCre, Nkx2.5Cre) causes morphological, metabolic and functional cardiac alterations that results in embryonic lethality [11, 12]. On the other hand, loss of function models based on conditional deletion of *Hif1a* gene in cardiac populations also causes several cardiac alterations during embryogenesis [13-15], indicating that a controlled balance in the oxygen levels and hypoxia signaling is required for proper heart development. However, there are important phenotypic discrepancies between the different published loss of function models. On one hand, the use of cardiomyocyte-specific Mlc2vCre line in combination with a *Hif1a* floxed mice does not affect embryonic survival but causes cardiac hypertrophy with reduced cardiac function in the adulthood, together with decreased glycolysis and ATP and lactate levels [14]. On the other hand, when one null *Hif1a* allele in germline is used in combination with a floxed allele and Mlc2vCre, mutant embryos show several cardiac alterations and increased cardiomyocyte proliferation, with associated embryonic lethality by E12.0 [15]. In contrast, a more recent study also uses a combination of null and floxed *Hif1a* alleles, but here recombination is driven by Nkx2.5Cre, an efficient Cre line expressed in cardiovascular progenitors contributing to the epicardium and endocardium in addition to cardiomyocytes [13]. In this case, *Hif1a* deletion seems to have opposite effects, with activation of cell stress pathways, such as p53 and ATF4, that inhibit cardiomyocyte proliferation and results in embryonic lethality by E15.5. Considering the importance of hypoxia pathway in early hematopoiesis, placentation and vascular development, the potential secondary effect of using *Hif1a*-null alleles on cardiac development cannot be ruled out when interpreting data from some of these mutants. Therefore, while sustained HIF1 signaling in the embryonic heart is detrimental for proper cardiac development, there is a lack of consensus about the impact of *Hif1a* loss during cardiogenesis and subsequent effects on cardiac function.

It has been widely demonstrated that HIF1 signaling is an important regulator of cellular metabolism, both in physiological and pathological contexts. In addition to glycolytic activation [16], HIF1 reduces mitochondrial metabolism by repressing pyruvate entry into the mitochondria [17] and by promoting COX4 isoform switch from COX4-1 to COX4-2 [18]. Moreover, HIF1 can also limit oxidative metabolism through an inhibitory role on mitochondrial biogenesis [19]. Several in vitro studies have demonstrated that the embryonic heart relies on glycolysis for energy supply [20, 21], in contrast with the adult heart that sustains most of ATP production through mitochondrial oxidation of fatty acids (FA) [22]. This metabolic switch is coincident with the change in oxygen levels after birth [23]. However, a recent report from our group has shown that metabolic shift towards fatty acid oxidation (FAO) occurs earlier during development at around E14.5, through a mechanism dependent of a decrease in HIF1 signaling in the embryonic myocardium [12]. Despite of the importance of glucose and FA as cardiac energy sources, amino acids can also be used as bioenergetics fuel. Hence, amino acids have the capacity to enter the Krebs cycle at different levels, a phenomenon known as anaplerosis, to replenish metabolic intermediates that warrant both NADH/FADH2 and building blocks production that enable the cells to continue growing under amino acid metabolism. The importance of amino acids as catabolic substrates has been described in several contexts, such as tumor growth [24], pulmonary hypertension [25] or limited oxygen supply conditions [26, 27]. However, the ability of the embryonic heart to catabolize amino acids remains unknown.

Here, we described that *Hif1a* loss during cardiogenesis blunts glycolysis and drives a compensatory metabolic adaptation based on transient activation of amino acid transport and catabolism associated with increased mitochondrial content to maintain energy production prior to the establishment of FA-based metabolism. Our results describe a new role of amino acid metabolism during heart development and open future research horizons towards studying the ability of the heart to use amino acids as an alternative energy fuel and biosynthetic precursor source in different pathophysiological contexts.

## METHODS

### Animal care and housing

*Hif1a*^*flox/flox*^ [28] mice were maintained on the C57BL/6 background and crossed with mice carrying *Nkx2.5Cre* recombinase [29] or *TnTCre* recombinase [30] in heterozygosity. *Hif1a*^*flox/flox*^ homozygous females were crossed with double heterozygous males and checked for plug formation. Mice were housed in SPF conditions at the CNIC Animal Facility. Welfare of animals used for experimental and other scientific purposes conformed to EU Directive 2010/63EU and Recommendation 2007/526/EC, enforced in Spanish law under Real Decreto 53/2013. Experiments with mice and embryos were approved by the CNIC Animal Experimentation Ethics Committee.

### Statistical analysis and data representation

For histological, immunohistochemical quantifications, electron microscopy and RT-qPCR, values were pooled for embryos with the same genotype from at least 3 independent litters and analyzed by the indicated statistical test using SPSS software (IBM; USA), with statistical significance assigned at P ≤0.05. Values were represented as mean ± SEM using GraphPad Prism (GraphPad; USA).

### Data access

RNASeq data: The accession number for the RNAseq data reported in this paper is GEO:

Proteomics data: The data set (raw files, protein databases, search parameters and results) is available in the PeptideAtlas repository (), which can be downloaded via ftp.peptideatlas.org (username: password:).

## RESULTS

### HIF1 signaling in cardiac progenitors is dispensable for cardiac development

HIF1α is expressed in the developing myocardium, with a temporal dynamics along midgestation [12, 13, 15]. We and others have described the heterogeneous regional distribution of HIF1 signaling with high HIF1α levels in the compact myocardium in contrast with low expression in the trabeculae [12, 13, 15]. HIF1 has been proposed to regulate cardiomyocyte proliferation during cardiogenesis. However, opposing results have been reported depending on the Cre line used for recombination and the utilization of one global null allele or two floxed alleles of *Hif1a* [13-15]. To investigate the role of HIF1 in heart development we generated a cardiac-specific loss of function model using floxed alleles of *Hif1a* gene in combination with the cardiac progenitor-specific Cre recombinase line *Nkx2.5*Cre (*Hif1a*^flox/flox^/*Nkx2.5*^Cre/+^, from here on *Hif1a*/*Nkx2.5*). Cre-mediated recombination was analyzed by agarose gel electrophoresis of cardiac and non-cardiac tissue at E12.5 (Fig. S1A). A 400bp product, corresponding to the processed *Hif1a* allele was only obtained in cardiac tissue in the presence of Cre recombinase activity, indicating that the deletion is specific in cardiac tissue and there is not ectopic recombination. To confirm the deletion efficiency, we analyzed by qPCR at E14.5 the expression of several genes involved in HIF1 pathway (Fig. S1B). Despite of an increase in *Hif1a* mRNA, probably associated with compensatory mechanisms, specific lack of amplification in the floxed region of the *Hif1a* gene showed efficient recombination. In addition, HIF1 target, *Phd3* showed decreased expression in the *Hif1a*/*Nkx2.5* mutants, while other HIF1 targets like *Vegfa* or additional elements of the hypoxia pathway like *Hif2a, Hif1b* or *Vhl* remain unchanged. We have also determined HIF1α protein distribution and abundance within the tissue by immunostaining in mutant and control littermates by E12.5 (Fig. S1C). *Hif1a* loss in the mutant embryos resulted in reduced HIF1α staining, with a displacement of the HIF1α channel intensity curve to lower fluorescence intensities (Fig. S1D). It is remarkable that the efficiency detected by qPCR is stronger than the one observed by immunostaining. This is probably because *Hif1a* mRNA after recombination is still stable to be translated into a protein, although this protein is not functional as it lacks the DNA-binding domain located in the N-terminal region of the floxed hif1a gene. The efficient deletion was further confirmed by Western-Blot of *Hif1a*-deficient hearts at E12.5 (Fig. 1A). The lower weight of HIF1α protein band confirms the presence of a truncated protein in the mutant embryos that is recognized by the HIF1 antibody raised against the C-terminal domain. *Hif1a*/*Nkx2.5* mutants were viable and recovered in the expected Mendelian proportions from E14.5 to weaning (Table 1). Histological analysis of control and *Hif1a*/*Nkx2.5* mutants at E12.5 (Fig. 1B), did not reveal differences between genotypes in terms of wall thickness (Fig. 1C) or sphericity (data not shown). Proliferation analysis by BrdU staining at E12.5 proved comparable proliferation index between control and *HIf1a*-deficient hearts (Fig. 1D). Similar results were obtained at E14.5, when *Hif1a*-deficient embryos did not show morphological alterations (Fig. S2A-B) nor differences between genotypes in terms of cell size (Fig. S2C) and proliferation by means of BrdU staining (Fig. S2D). These results indicate that the lack of *Hif1a* in Nkx2.5 cardiac progenitors does not influence cardiomyocyte proliferation and suggest that HIF1 signaling is dispensable for proper cardiac development.

**Table 1.** Embryo recovery analysis of *Hif1a*/*Nkx2.5* mutants. For each stage, table shows the number of mutant embryos recovered, the total number of embryos/pups collected and the number of litters analyzed. The percentage of recovered mutants and the expected recovery percentage (25% in all cases) were compared by the Wilcoxon signed rank test.

| Stage | *Hif1a<sup>f/f</sup>/Nkx2.5<sup>Cre/+</sup>* | Total | Litters | Observed % | Expected % | p-value |
| --- | --- | --- | --- | --- | --- | --- |
| E14.5 | 20 | 90 | 13 | 24.4±4.9 | 25 | 0.905 |
| Weaning | 10 | 38 | 6 | 28.6±5.8 | 25 | 0.787 |

**Figure 1.**
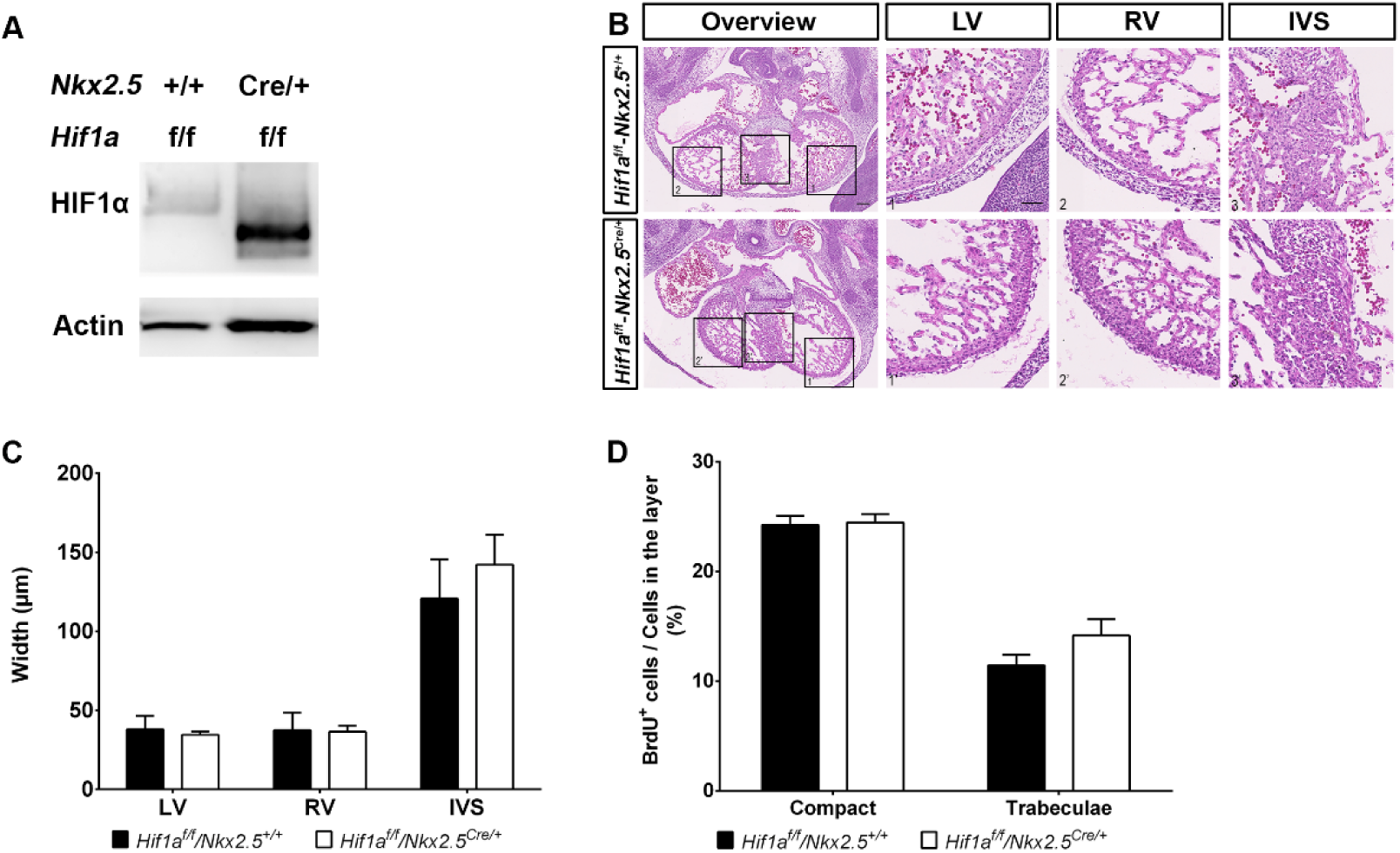
Embryonic phenotype of *Hif1a*-deficient embryos at E12.5. **A)** Representative immunoblot against HIF1α (up) and α-Actin (down) in heart lysates of control (*Hif1a*^f/f^/*Nkx2.5*^+/+^) and *Hif1a*/*Nkx2.5* (*Hif1a*^f/f^/*Nkx2.5*^Cre/+^) embryos at E12.5. **B)** E12.5 control (up) and mutant (down) embryos stained with hematoxylin and eosin (HE). Scale bars represent 100µm (overview) and 20µm (insets). **C)** HE quantification of ventricular walls and inter-ventricular septum width in E12.5 control (black bars) and mutant (white bars) embryos. **D)** Quantification of BrdU immunostaining, represented as percentage of BrdU^+^ cells in the compact myocardium and trabeculae of E12.5 control (black) and *Hif1a/Nkx2.5* mutant (white) embryos. In all graphs, bars represent mean ± SEM (n=3), Student’s t test, * p value<0.05.

### Cardiac deletion of *Hif1a* prevents the expression of glycolytic enzymes in the compact myocardium

To investigate adaptive mechanisms operating upon *Hif1a* loss to allow normal cardiac development, we performed massive expression analysis by RNASeq of E12.5 ventricular tissue from control and *Hif1a*/*Nkx2.5* mutant embryos. Subsequent bioinformatics analysis identified 14406 protein-coding genes being expressed. Among them, 201 genes showed differential expression between control and mutant hearts: 118 were downregulated and 83 were upregulated (Table S1). Gene enrichment analysis revealed that the absence of HIF1 produced significant alterations in metabolic processes, such as nucleotide/nucleoside and monocarboxylic acid metabolism, amino acid and, especially, carbohydrate metabolism (Table S2).

We have previously described the existence of metabolic territories in the embryonic myocardium with an enhanced glycolytic signature in the compact myocardium by E12.5 [12]. Here we show that the main affected functions in the *Hif1a*/*Nkx2.5* mutant hearts were related with glucose catabolic processes and transport (Fig. 2A). Moreover, glucose transporter 1 (GLUT1) protein levels were significantly reduced in the compact myocardium of the *Hif1a*/*Nkx2.5* mutant embryos by E12.5 (Fig. 2B). The inhibition of the glycolytic program in the *Hif1a*/*Nkx2.5* single mutant was further confirmed by mRNA expression analysis of the critical enzymes *Glut1, Pdk1* and *Ldha* by *in situ* hybridization at E12.5 and E14.5 in control and *Hif1a/Nkx2.5* mutants (Fig. 2C). Results showed strong inhibition of glycolytic gene expression in the compact myocardium of *Hif1a*-deficient hearts at both stages. This sustained glycolytic inhibition at E14.5 was validated by qPCR (Fig. 2D), including the downregulation of the monocarboxylic acid transporter *Slc16a3*, responsible of monocarboxylic acid transport, such as lactate, across the plasma membrane. NKX2.5 cardiac progenitors contribute to different cardiac layers including myocardium, epicardium and endocardium. To determine if glycolytic inhibition was associated with *Hif1a* loss in other cardiac layers, we specifically deleted *Hif1a* in cardiomyocytes using *cTnT-*Cre [30](*Hif1a*^flox/flox^/*cTnT*^Cre/+^, hereon *Hif1a*/*cTnT*). *Hif1a*/*cTnT* mutants also showed reduced HIF1α levels by E14.5 (Fig. S3A) compared with control littermates. Similar to *Hif1a*/*Nkx2.5* mutants, deletion of *Hif1a* in cardiomyocytes did not cause cardiac developmental defects by E14.5 (Fig. S3B) even in the absence of effective glycolysis, as confirmed by *in situ* hybridization (Fig. S3C) and RT-qPCR (Fig.S3D) of key glycolytic pathway members. Taken together, these results confirm our previous findings that HIF1 signaling controls the expression of glycolytic genes in the embryonic heart and indicate that an active glycolytic program in the compact myocardium is not essential for normal cardiogenesis.

**Figure 2.**
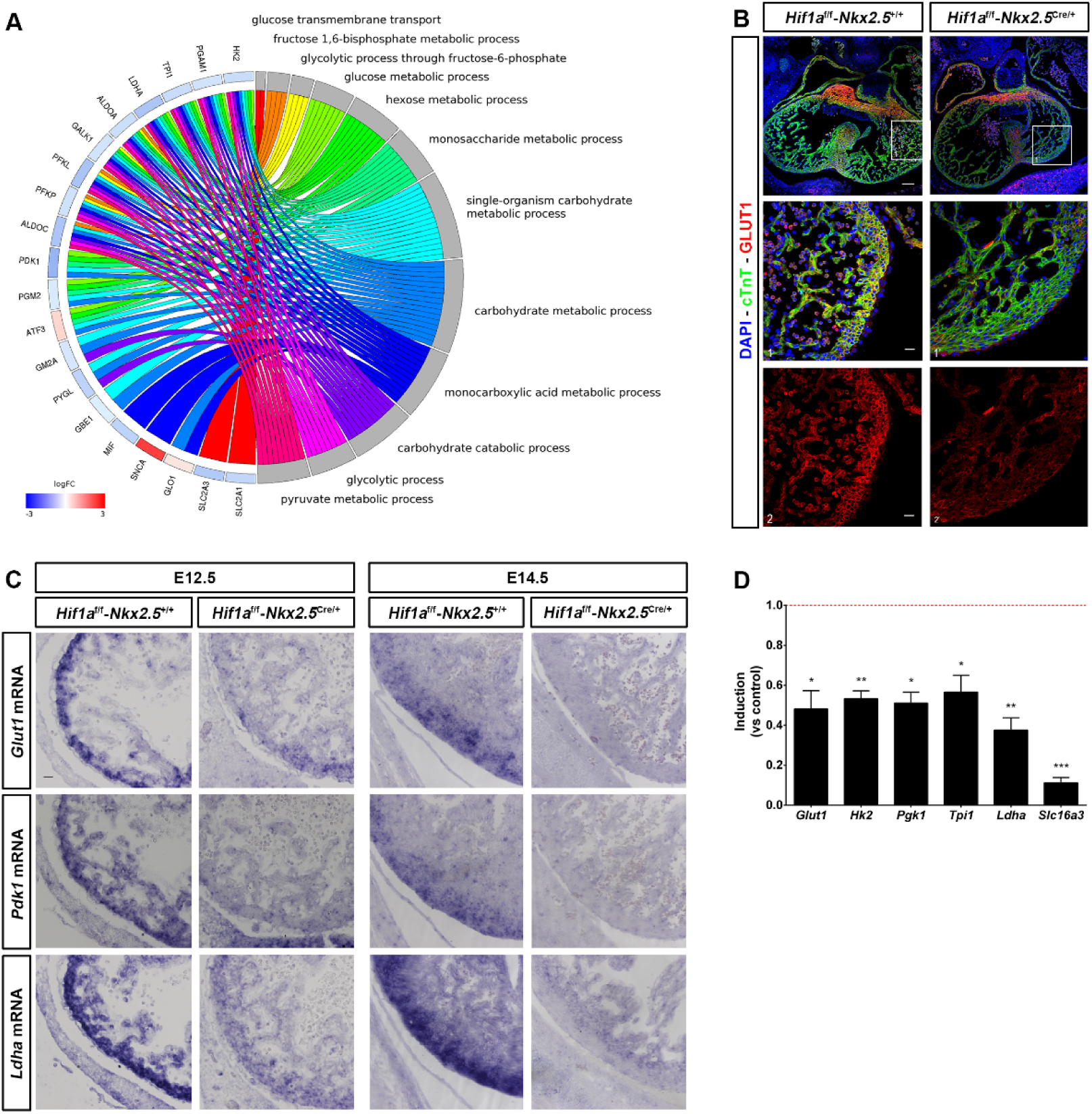
Glycolytic metabolism alterations in *Hif1a*/*Nkx2.5* embryos. **A)** Differentially expressed genes extracted from *Hif1a*/*Nkx2.5* RNASeq analysis at E12.5 and associated enriched GO terms for the category “Carbohydrate metabolism” according to the Gorilla database. Expression changes are shown as induction in mutant (*Hif1a*^f/f^/*Nkx2.5*^Cre/+^) relative to control embryos (*Hif1a*^f/f^/*Nkx2.5*^+/+^) and color coded by logarithmic fold change (log FC). **B)** GLUT1 immunofluorescence on E12.5 heart sections of control (left) and *Hif1a/Nkx2.5* mutants (right). Nuclei shown in blue, Troponin T in green and GLUT1 in red. Insets show left ventricle. Scale bars, 100µm and 20µm in insets. **C)** In situ hybridization of *Glut1* (top), *Pdk1* (middle) and *Ldha* (bottom) in control and *Hif1a-*mutant right ventricle at E12.5 (left) and E14.5 (right). Scale bar, 20µm. **D)** RT-qPCR analysis of glycolytic genes in E14.5 *Hif1a*-mutant ventricles. Bars (mean ± SEM, n=3) represent fold induction relative to baseline expression in littermate controls (red line). Student’s t test. *p value<0.05; **0.005<p value<0.01, ***p value<0.005.

### Lack of cardiac *Hif1a* promotes precocious mitochondrial maturation without affecting lipid metabolism

We have previously reported that sustained HIF1 signaling in the embryonic myocardium results in severe alterations of mitochondrial amount and function [12]. To evaluate the bioenergetics adaptations in response to lack of cardiac HIF1 activation we investigated mitochondrial network and activity in *Hif1a*/*NKx2.5* mutants. Analysis and quantification of mitochondrial content by transmission electron microscopy (TEM) at E14.5 (Fig. 3A, B) revealed comparable mitochondrial amount between control and *Hif1a*-defficient embryos. Similarly, Cytochrome Oxidase 4 (COX4) activity did not show differences between genotypes at E14.5 (Fig. 3C), with higher mitochondrial activity in the trabeculae of control and *Hif1a*-deficient hearts as previously reported [12]. Despite the absence of mitochondrial alterations, the onset of FA catabolism could be affected in *Hif1a*-null hearts. To exclude this possibility, we analyzed intracellular lipid accumulation by TEM in ventricular tissue of *Hif1a*/*Nkx2.5* mutants and control embryos by E14.5. Quantification of the lipid droplet area (Fig. 3D) indicated no significant differences in terms of accumulation of intracellular lipids in *Hif1a*-deficient embryos compared with control littermates.

**Figure 3.**
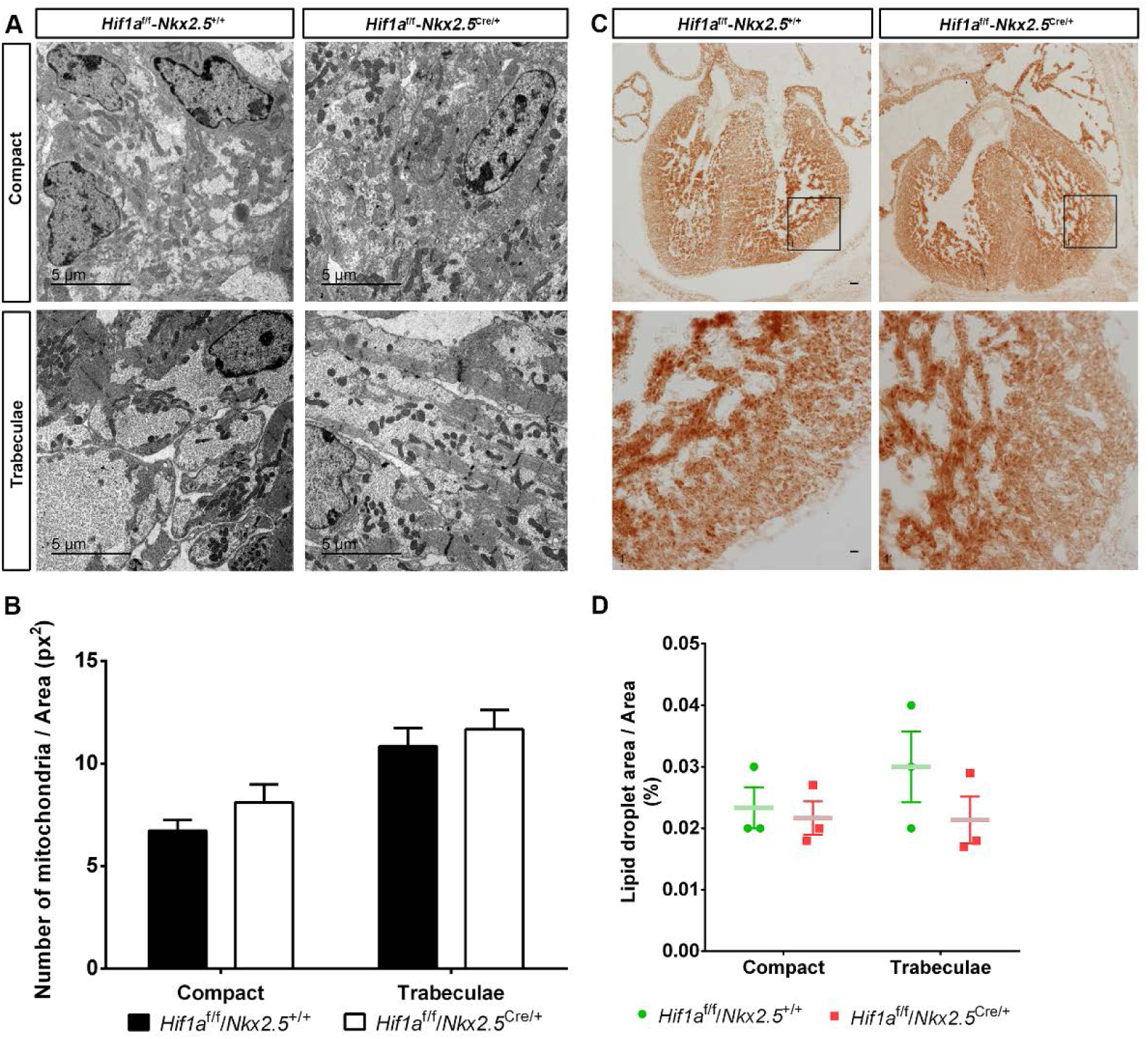
Mitochondrial metabolism in *Hif1a*/*Nkx2.5* mutants at E14.5. **A)** Transmission electron micrographs of ventricular tissue from a representative E14.5 control embryo (*Hif1a*^*f/f*^/*Nkx2.5*^*+/+*^, left) and a mutant littermate (*Hif1a*^*f/f*^/*Nkx2.5*^*Cre/+*^, right), showing compact myocardium (top) and trabeculae (bottom). **B)** Quantification of total mitochondria in electron micrographs from E14.5 controls (black bars) and mutants (white bars). Results are expressed as number of mitochondria per tissue area (px^2^). **C)** Representative unfixed cardiac sections from E14.5 control embryos (left) and mutant littermates (right) stained for cytochrome c oxidase subunit 4 (COX4) activity. Scale bars, 100µm (upper panels) and 20µm (lower panels). **D)** Percentage of lipid droplet area relative to tissue area in cardiac electron micrographs of E14.5 control embryos (green dots) and *Hif1a*-mutant littermates (red squares). In all graphs, bars represent mean ± SEM (n=3), Student’s t test.

These results indicate that *Hif1a* deletion does not preclude the establishment of an active mitochondrial network and that the embryonic FA metabolism is conserved in cardiac *Hif1a* mutants by E14.5.

Since HIF1α levels are stronger at gestational stages earlier than E14.5 [12], we decided to determine the effect of *Hif1a* loss on the mitochondrial network by E12.5, when endogenous HIF1α is strongly expressed in the compact myocardium. Analysis and quantification of ventricular ultrastructure by TEM at E12.5 (Fig. 4A-B) indicated increased mitochondrial content in *Hif1a*-deficient embryos compared with control littermates, despite of the mitochondrial enrichment in the trabeculae was still present in the mutant embryos.

**Figure 4.**
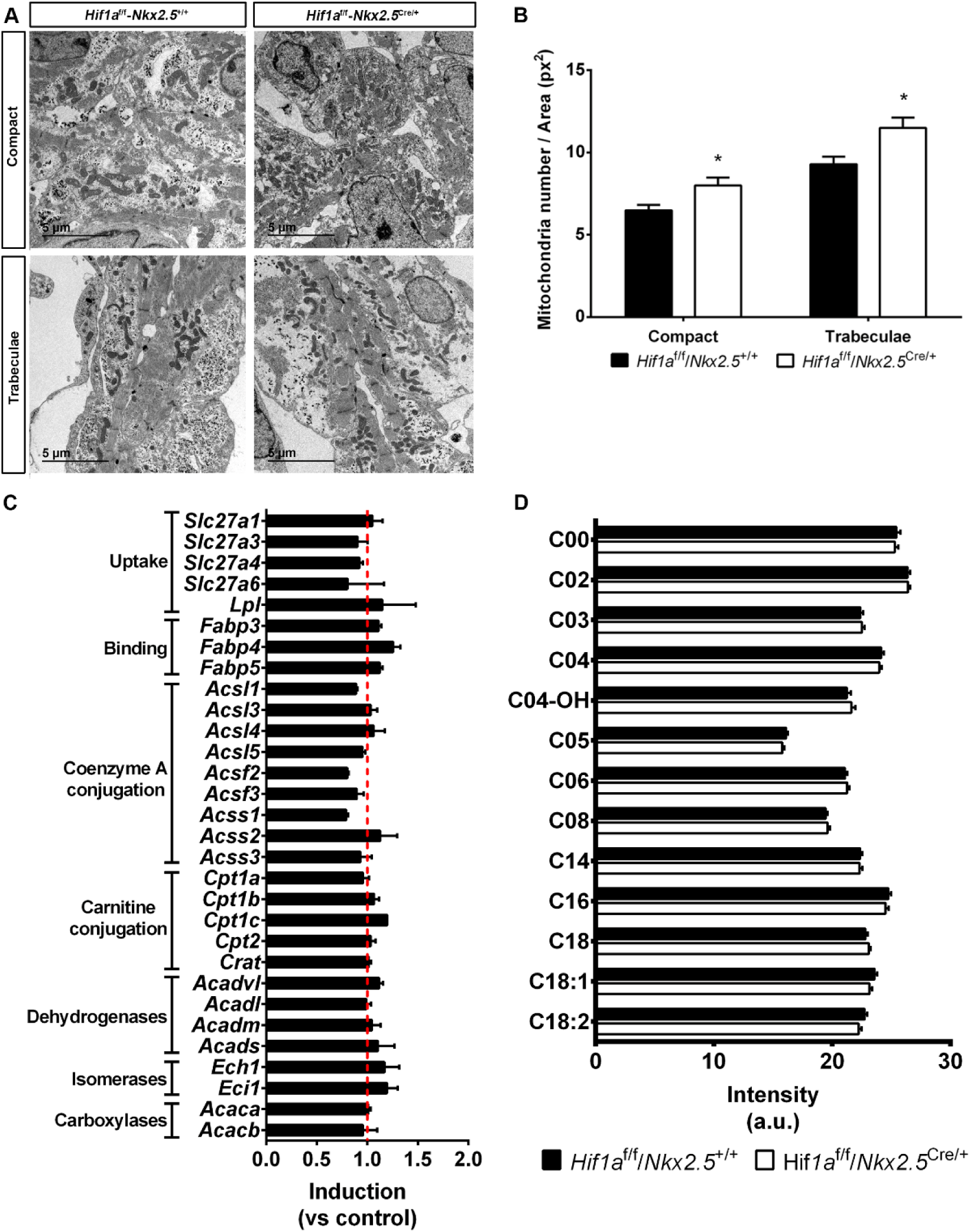
Mitochondrial metabolism in *Hif1a*/*Nkx2.5* mutants at E12.5. **A)** Transmission electron micrographs of ventricular tissue from a representative E14.5 control embryo (*Hif1a*^*f/f*^/*Nkx2.5*^*+/+*^, left) and a mutant littermate (*Hif1a*^*f/f*^/*Nkx2.5*^*Cre/+*^, right), showing compact myocardium (top) and trabeculae (bottom). **B)** Quantification of total mitochondria in electron micrographs from E14.5 controls (black bars) and mutants (white bars). Results are expressed as number of mitochondria per tissue area (px^2^). Bars represent mean ± SEM (n=3) **C)** Fold change gene expression determined by RNASeq of genes involved in fatty acid uptake and catabolism in *Hif1a*/*Nkx2.5* mutants. Red line represents baseline expression in control littermates. Bars represent mean ± SEM (n=2) **D)** Quantification of fatty acid-conjugated carnitines by metabolomic untargeted profiling using MS/MS in control (black bars) and *Hif1a*/*Nkx2.5* (white bars) hearts at E12.5. Bars represent mean ± SEM (n=13). In all graphs, bars represent mean ± SEM (n=3), Student’s t test, \**P*<0.05.

As mature cardiomyocytes rely on FAO for ATP production and cardiac performance, one possible metabolic adaptation of *Hif1a* deficient hearts associated with the increased mitochondrial content could be an early utilization of FA to provide sufficient ATP levels in the absence of effective glycolysis. However, genes involved in lipid catabolism did not show altered expression between genotypes at E12.5 in the RNASeq analysis (Fig. 4C). Moreover, carnitine and acyl-carnitine metabolite MS profiling did not reveal significant alterations in *Hif1a*/*Nkx2.5* mutants (Fig. 4D).

These results demonstrate that reduced HIF1 signaling promotes a premature development of a cardiac mitochondrial network and suggest the activation of compensatory mechanisms other than FAO activation upon glycolytic inhibition in *Hif1a* mutant embryos

### Amino acid metabolism is transiently enhanced in the embryonic heart in the absence of effective glycolysis in Hif1α mutant mice

The increased mitochondrial content detected in *Hif1a*-deficient embryos by E12.5 was confirmed by significant enriched expression of genes related to oxidative phosphorylation determined by Gene Set Enrichment Analysis (GSEA) (Fig. 5A). To better understand the adjustment in cardiac metabolism upon *Hif1a* deletion, we further investigated the GO analysis performed at E12.5 and found altered processes related to amino acid metabolism in *Hif1a/Nkx2.5* mutants. More specifically, GO analysis showed alterations in processes related to amino acid transport and metabolism as well as Ala, Leu, Val, Ile, Asn, Asp, Ser, and Gly biosynthesis (Fig. 5B, Table S2, Table S3). These results lead us to hypothesize about the activation of a metabolic reprogramming towards amino acid oxidative catabolism of embryonic cardiomyocytes in the absence of effective glycolysis associated to *Hif1a* deletion.

**Figure 5.**
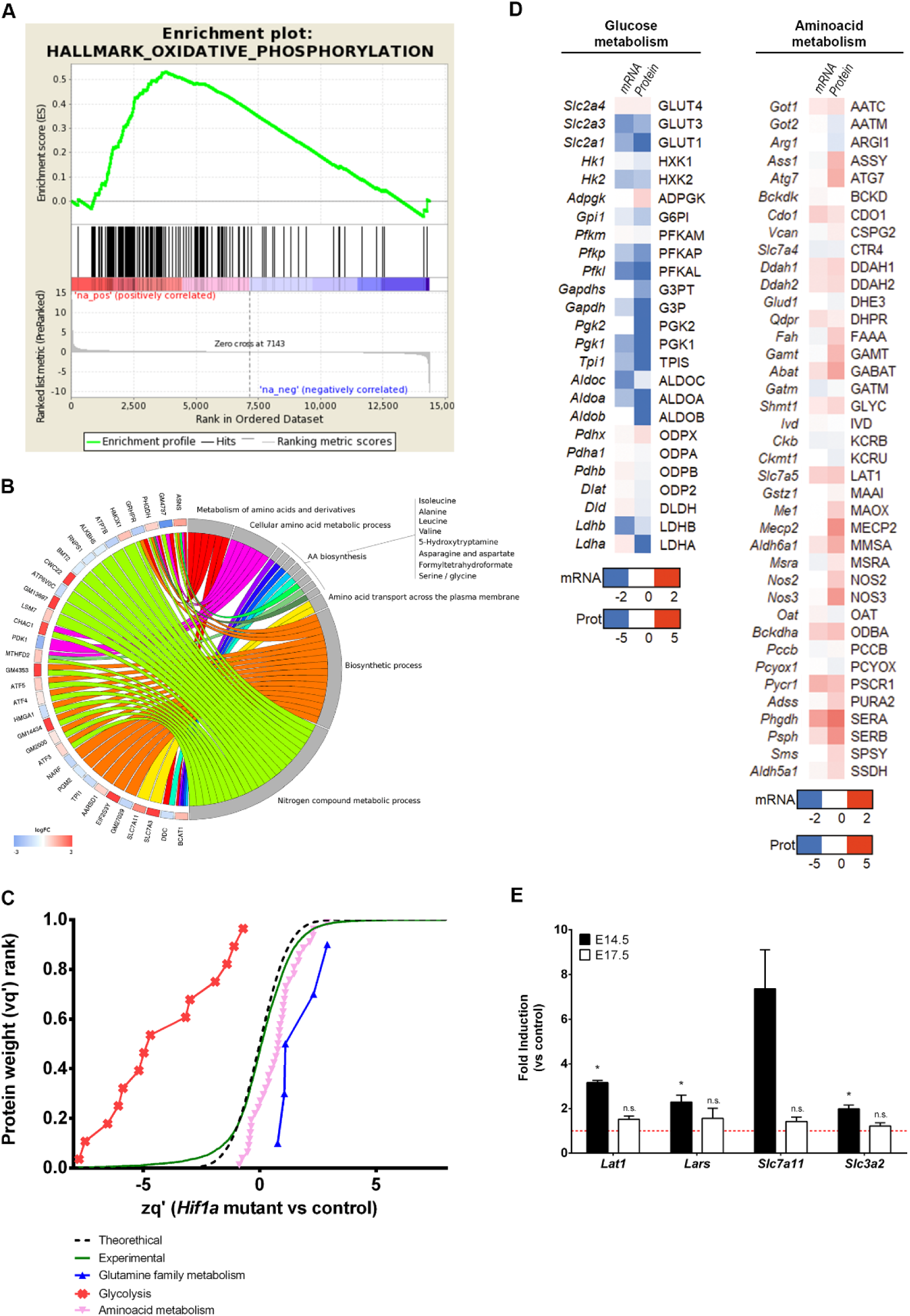
Metabolic adaptations in *Hif1a*-deficient hearts at E12.5. **A)** GSEA enrichment analysis for Oxidative Phosphorylation term in Hallmark database. “na_pos” in red corresponds to *Hif1a*/*Nkx2.5* mutants (*Hif1a*^*f/f*^/*Nkx2.5*^*Cre/+*^) and “na_neg” in blue correspond to control (*Hif1a*^*f/f*^/*Nkx2*^*r+/+*^) littermates. **B)** Differentially expressed genes extracted from *Hif1a*/*Nkx2.5* RNASeq analysis at E12.5 and associated enriched GO terms for the category “Amino acid metabolism” according to the Gorilla database. Expression changes are shown as induction in mutant (*Hif1a*^f/f^/*Nkx2.5*^Cre/+^) relative to control embryos (*Hif1a*^f/f^/*Nkx2.5*^+/+^) and color coded by logarithmic fold change (log FC). **C)** Representation of protein statistical weights (wq’) grouped by functional categories (FDR<1%, n=6) versus protein abundance in *Hif1a*-deficient hearts relative to control embryos (zq’) at E12.5, as determined by MS/MS proteomics. A displacement right from the experimental curve indicates increased pathway in mutant embryos. **D)** Heatmap representation of mRNA (quantified by RNASeq) and Protein (quantified by MS/MS) of components of glucose (left) and amino acid (right) metabolic pathways. Color code indicated in the legend is calculated as the value found in *Hif1a*/*Nkx2.5* mutants relative to control littermates. **E)** RT-qPCR analysis of amino acid transporter gene expression in E14.5 (black bars) and E17.5 (white bars) *Hif1a*-mutant ventricular tissue. Bars (mean ± SEM, n=2 E14.5 and n=3 for E17.5) represent fold induction relative to baseline expression in littermate controls (red line). Student’s t test, * p value<0.05, n.s. non-significant.

To validate this hypothesis, we performed global proteomic analysis in *Hif1a*/*Nkx2.5* embryos and control littermates by E12.5. System Biology analysis of the proteome quantification (Fig. 5C) revealed significant increase in proteins related to amino acid metabolism and, specifically, glutamine family metabolism. In addition, the observed decrease in proteins related to glycolysis confirmed the inhibited expression of glycolytic enzymes upon *Hif1a* deletion (Fig. 2). Moreover, the quantification of both gene expression by RNASeq and protein abundance by MS/MS proteomics display parallel results, showing inhibited glycolysis (Fig. 5D, left) and increased amino acid catabolism (Fig. 5D, right). Specifically, *Hif1a*/*Nkx2.5* mutants showed increased expression and protein levels of genes potentially contributing to anaplerosis of amino acids into Krebs’ Cycle. These contributions (Fig. S4) included aromatic amino acids (Phe, Tyr), polar amino acids (Asn, Asp, Gln, Glu, Ser, and Cys), Pro, Gly and branched-chain amino acids (Val, Leu, and Ile). Enzymes involved in the Urea Cycle were also upregulated, what could result in increased contribution of Arg to Krebs’ Cycle by the generation of fumarate. To determine whether this amino acid signature was maintained over time in *Hif1a*-deficient hearts, we analyzed the expression of amino acid transporters by RT-qPCR at E14.5 and E17.5 (Fig 5E). The results show increased gene expression levels of several transporters, such as *Lat1* or *Slc7a5* that form heterodimers with *Slc3a2*, also upregulated, (transporter of branched and aromatic amino acids like Met, Val, Leu, Ile, Trp, Phe, Tyr and His) [31] [32] and *Slc7a11* (transporter of Cys) [33].In addition, the leucyl-tRNA synthetases *Lars* and *Lars2*, were still upregulated by E14.5 in *Hif1a*-deficient hearts, but reach control-like expression levels by E17.5.

These results support the transient upregulation of amino acid catabolism and anaplerosis upon *Hif1a* loss, uncovering the metabolic flexibility of the embryonic heart to adapt to different substrates for energy supply. Moreover, our data subscribe the idea that *Hif1a*-deficient hearts are still able to switch their metabolism towards FAO from E14.5.

### Developmental loss of *Hif1a* does not influence adult cardiac function or morphology

Albeit deletion of *Hif1a* did not hamper cardiac development, we wondered whether mutant mice might develop heart alterations in the adulthood. Histology analysis by Hematoxylin-Eosin (HE) staining at five months of age did not indicate evident changes in cardiac morphology of *Hif1a*-deficient hearts relative to control animals (Fig. 6A). Moreover, Mason’s Trichrome staining analysis excluded the presence of fibrotic areas in any of the genotypes (Fig. 6B). Nevertheless, the lack of macroscopic malformations does not rule out that cardiac performance could be affected. To determine if embryonic deletion of *Hif1a* influences cardiac function during the adulthood, we performed echocardiography to five months-old control and mutant mice. Both 2D and M mode analysis (Fig. 6C) and the quantification of several echocardiographic parameters (Fig. S5A-D) confirmed the absence of anatomical alterations. Furthermore, conserved cardiac function in *Hif1a/Nkx2.5* mutant versus control mice was demonstrated by means of Ejection Fraction (EF) and Fractional Shortening (FS) (Fig. 6D). On the other hand, electrocardiographic analysis showed normal PR and QRS segment length in *Hif1a*/*Nkx2.5* mice (data not shown), ruling out the existence of conduction or coupling defects. These results indicate that active HIF1 signaling is not required for normal cardiac function in the adult heart.

**Figure 6.**
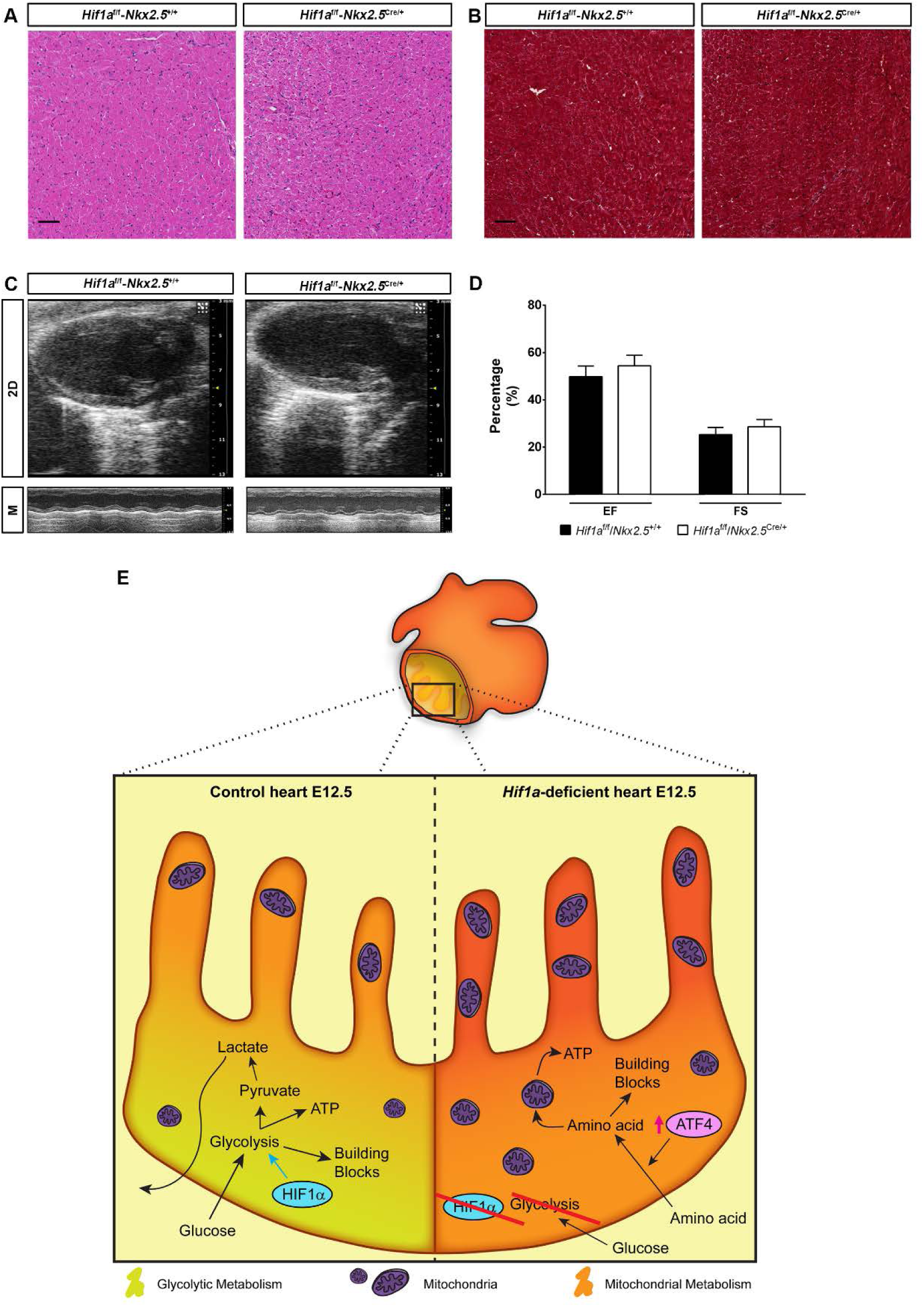
Cardiac morphology and function in adult *Hif1a*/*Nkx2.5* mutants. HE (**A**) and Mason’s Trichrome (**B**) stainings of the left ventricle from 5 months-old control (*Hif1a*^*f/f*^/*Nkx2*^*r+/+*^) and *Hif1a*/*Nkx2.5* mutants (*Hif1a*^*f/f*^/*Nkx2.5*^*Cre/+*^). Scale bar represents 50µm. **C**) Echocardiography analysis of 5 months-old control and *Hif1a*-deficient mutants in 2D mode (up) and M mode (down). **D)** Quantification of Ejection Fraction (EF) and Fractional Shortening (FS) in control (black bars, n=9) and *Hif1a*/*Nkx2.5* mutants (white bars, n=11) by 5 months of age. Bars represent mean ± SEM, Student’s t test. **E)** Model representing the embryonic myocardium by E12.5. Compact myocardium is mainly glycolytic (yellowish), by the action of HIF1 signaling, while trabeculae rely more on mitochondrial metabolism (orange) in control embryos (left). In *Hif1a*-mutants (right), the whole myocardium relies on mitochondrial metabolism and present higher mitochondrial content, using amino acids as energy source possibly through the compensatory activation of ATF4 activation.

In summary, our results demonstrate that HIF1 signaling in NKX2.5 cardiac progenitors is dispensable for proper heart development and that the absence of *Hif1a* triggers a cardiac metabolic reprogramming, enhancing amino acid catabolism to ensure sufficient ATP and biosynthetic precursors to sustain cardiac growth and function in the absence of effective glycolysis (Fig. 6E). Importantly these adaptations might be relevant in the adulthood under pathological scenarios associated with increased HIF signaling like pulmonary hypertension or cardiomyopathy towards the development of new drugs against new metabolic targets.

## DISCUSSION

Here, we describe that *Hif1a* loss in NKX2.5 cardiovascular progenitors causes glycolytic program inhibition in the compact myocardium by E12.5, without compromising normal cardiac development and embryonic viability. Our results show that upon *Hif1a* deletion, the embryonic myocardium exhibits the ability to activate metabolic programs oriented to amino acid catabolism, together with a premature increase in the mitochondrial content by E12.5. Taken together, our findings point out the metabolic versatility of the embryonic heart and conciliate the discrepancies from previous deletion models of *Hif1a* in cardiovascular progenitors.

### Integration with previous *Hif1a* deletion model in the embryonic heart

As outlined in the introduction, there is a lack of consensus between previous reports on cardiac embryonic mouse models of *Hif1a* loss. On the one hand, the use of Mlc2vCre, a mild and not homogenously expressed driver within the myocardium as *Nkx2.5* [34], might lead to heterogeneous recombination pattern and hence not complete abrogation of HIF1 signaling in cardiomyocytes that could cause the adult phenotype described by Huang and collaborators [14]. Thus, the use of different Cre drivers and its differences in terms of recombination pattern and efficiency keeps off any kind of comparison between genetic models. On the other hand, previous reports from Perriard and Zambon/Evans’ group [13, 15], despite of the Cre driver applied, use a null allele of *Hif1a* in combination with a floxed allele. The *Hif1a* haploinsufficiency of these models outside the heart NKX2.5-territories might cause extracardiac affections, such as vascular or placental defects, that could significantly influence the described phenotype. In this regard, the exhaustive characterization by Guimaraes-Camboa et al. at gene expression level extensively overlaps with our RNASeq data, except in terms of stress and apoptotic pathways, upregulated only in their model combining null and floxed alleles.

Moreover, we have also analyzed the *Hif1a* null/floxed mice in NKX2.5 progenitors in parallel with the double floxed model described here and, even though the glycolytic inhibition by E14.5 is comparable in both models (data not shown) only the null/floxed mice exhibited embryonic lethality (5% retrieved versus 25% expected, p value 0.0029, n=7 litters). This observation, together with our results, support the notion that *Hif1a* is dispensable for cardiac development, while an extensive comparison, in terms of gene expression in placental and vascular embryonic tissue, between the double floxed and the null/floxed models would be necessary to exclude extracardiac influences of *Hif1a* deficiency impacting on heart development as reported by Guimaraes-Camboa and colleagues.

### Amino acid catabolism and metabolic versatility of the embryonic heart

A key finding of our investigation is the fact that the embryonic myocardium is able to upregulate alternative metabolic pathways (amino acids catabolism), and to promote mitochondrial biogenesis that could support the ATP demand upon glycolytic inhibition subsequent to *Hif1a* loss. The use of amino acids as cardiac metabolic fuel has been proposed mainly in oxygen-deprived scenarios [26, 27]. Amino acids provide, by deamination, carbon skeletons that can be converted into pyruvate, alpha-ketoglutarate, succinyl-CoA, fumarate, oxalacetate, Acetyl-CoA and Acetoacetyl-CoA, all of them metabolites that can be incorporated into the Krebs cycle [35, 36]. As detailed above, our *Hif1a*-defficient model upregulates both, at the transcriptional and protein level, a variety of amino acid transporters, biosynthetic and catabolic enzymes, that can replenish the Krebs cycle’s upon glucose deprivation. Moreover, the fact that this upregulation is accompanied by a premature increase in mitochondrial content suggests that the embryonic heart, in the absence of *Hif1a*, readapt its metabolism to maintain enough ATP levels and building blocks, without compromising the normal protein synthesis required for myocardium development and embryo viability.

Interestingly this adaptation is transient and reversible, as seen by the control-like levels of amino acid transporters transcripts found at later stages by E17.5, without precluding the embryonic metabolic switch towards FAO as previously described [12]. This shows that the embryonic myocardium has the plasticity to modulate its metabolism to adapt to the energetic demand and nutrient availability. Moreover, in addition to this interesting role of amino acid catabolism activation in the embryonic context, the use of amino acids as an alternative energy source could be an interesting option to achieve cardioprotection and recovery after cardiac injury. In this regard, some of the enzymes upregulated in our massive screenings in *Hif1a*-deficient hearts are involved in Ser biosynthesis and one-carbon cycle, including *Phgdh, Psph* and *Shmt1*. These pathways have been recently described to increase glutathione levels and protect the heart against oxidative stress [37], also in a context of myocardial hypertrophy [38].

### Origin of catabolized amino acids in *Hif1a***-deficient hearts**

One interesting open aspect of the metabolic adaptation exhibited by the *Hif1a*-deficient hearts is what would be the source of amino acids supply during glycolytic inhibition upon *Hif1a* loss. In this regard, two potential sources could be considered. First, protein-forming amino acids could be recycled through autophagy. This hypothesis is reasonable considering the context of the embryonic heart, where protein turnover, specially transcription factors, happens fast and at a high rate [39]. Moreover, a positive nitrogen balance has been reported in both adult rat and human hearts, indicative of rapid turnover of tissue proteins [40]. Interestingly, our *Hif1a* mutant embryos showed increased transcription of *p62*. P62 is a cargo-recognizing protein involved in autophagic degradation of cellular proteins [41]. While an extensive characterization analyzing autophagy, pro and anti-autophagic signaling pathways and protein labelling and turnover would be needed to further investigate this hypothesis, the fact that autophagy could be involved in this metabolic adaptation suggest an exciting link between cardiac metabolism, hypoxia and autophagy.

Another possible source of amino acids in *Hif1a*-deficient hearts is the fetal circulation. Even though an extensive characterization of fetal blood nutrient content over gestation has not been reported, the transcriptional increase in several membrane amino acid transporters observed in our mutants suggest that *Hif1a*-deficient cardiomyocytes could be obtaining them directly from embryonic circulation. Interestingly, cardiac amino acids uptake in human subjects infused intravenously with protein hydrolysate increases by 245% [26], showing that the heart can respond to blood amino acids levels. Moreover, the regulation of amino acid transporters expression in the placenta is essential for maintaining high levels of amino acids in the fetal blood to sustain embryo growth [42]. In this regard, an increased cardiac uptake of amino acids in the *Hif1a*-deficient embryo could result in increased amino acids supply through the placenta that might response to some secreted cues in the absence of cardiac HIF1 signaling.

### Molecular determinants of amino acid catabolism activation

Amino acids metabolism and transport is tightly regulated through several pathways, including mTOR, GCN2 (general control non-derepressable 2) and G-protein-coupled receptors, among others [43]. ATF4 is a transcriptional regulator that activates the expression of genes involved in amino acid transport and metabolism [44] and also respond to nutrient and metabolic stress in low oxygen conditions [45, 46] *Atf4* gene expression is positively regulated in our *Hif1a* deletion model at the transcriptional level. In addition, higher levels of ATF4 protein were found in the mutant embryos at E12.5, E14.5 and E17.5, suggesting that ATF4 pathway activation could be regulating, at least partially, the increase in the amino acid catabolic program shown in these embryos. Interestingly, *Hif1a* deletion model by Guimaraes-Camboa and collaborators [13] also shows increased ATF4 signaling in *Hif1a*-deficient hearts, suggesting that ATF4 could be in fact one of the main regulators of the described metabolic adaptation upon loss of effective glycolysis in the *Hif1a-*deficient hearts. Additionally, since ATF4 is upregulated in both animal models, and considering the lack of lethality of our floxed/floxed mice, the upregulation of ATF4 stress pathway does not seem to be responsible of the embryonic lethality reported by Zambon’s/Evan’s groups.

In parallel to ATF4 activation, HIF1 loss could cause a compensatory transient increase in HIF2α isoform that could be related with the increased amino acid catabolism transcriptional program observed in our *Hif1a*-defficient model. HIF2 has been described to activate cMYC in cancer cells [47]. Interestingly, cMYC promotes amino acid metabolism, including the expression of *Slc7a5* [24] or LAT1, involved in the transport of large neutral amino acids, such as Arg, Leu or Tyr. HIF2α has been also involved in the direct expression of LAT1 by binding to the proximal promoter of the gene in WT8 renal clear cell carcinoma cell line, as well as in lung and liver cells [48]. Hence, it would be of interest to investigate whether HIF2α could be an additional potent inductor of amino acid catabolic pathways, setting the basis for further research in metabolic adaptations by HIF2 in the absence of active HIF1 signaling.

In summary, our data show a link between *Hif1a* loss and cardiac metabolic reprogramming during embryo development. The results shown here highlight the metabolic versatility of the embryonic heart and open new research avenues to investigate how amino acid metabolism could play a relevant role in adaptive responses to stress conditions in the heart.

## Supporting information

This file contains supplemental methods, figures, tables, legends and references to the manuscript by Menendez-Montes et al,

## NON-STANDARD ABBREVIATIONS AND ACRONYMS”

FA: Fatty Acids
FAO: Fatty Acid Oxidation
GSEA: Gene Set Enrichment Analysis
GO: Gene Ontology
HE: Hematoxylin-Eosin

## ACKNOWLEDGMENTS

Authors thank Lorena Flores for echocardiography technical assistance, Raquel Baeza for animal housing and handling and to CNIC Microscopy, Genomics and Histology Core Facilities for technical assistance. We thank Luke Szweda, Hesham Sadek, Julian Aragones and Miguel Torres for critical revision and discussion of the manuscript.

