## Supplementary material for "Metabolic reprogramming from glycolysis to amino acid utilization in cardiac HIF1α deficient mice": This file contains supplemental methods, figures, tables, legends and references to the manuscript by Menendez-Montes et al,

**SUPLEMENTAL MATERIAL**

**DETAILED METHODS**

**Genotyping**

Genotyping was performed as previously described (*1*), using the following primers (Sigma Aldrich; USA) for *Hif1a floxed* alleles: 5’ CGTGTGAGAAAACTTCTGGATG 3’ and 5’ AAAAGTATTGTGTTGGGGCAGT 3’. For *Hif1a* null allele detections the primers were: 5’ GCCCATGGTAAGAGAGTAGGTGGG 3’ and 5’ 5’ AAAAGTATTGTGTTGGGGCAGT 3’. For Cre alleles genotyping, the following primers were used: *Nkx2.5*: 5’ GCCCTGTCCCTCAGATTTCACACC 3’, 5’ GCGCACTCACTTTAATGGGAAGAG 3’ and 5’ GATGACTCTGGTCAGAGATACCTG 3’ and *cTnT*: 5’ TACTCAAGAACTACGGGCTGC 3’ and 5’ GCACTCCAGCTTGGTTCCCGA 3’.

**Embryo extraction**

Embryos were extracted after pregnant female euthanasia by $CO_2$ inhalation and head and liver were removed. Embryos were then fixed overnight at 4ºC in 4%PFA solution (RT15710, Electron Microscopy Sciences; USA). After fixation, embryos were dehydrated, embedded in paraffin and sectioned at 5µm for immunostaining and histological purposes and at 10µm for in situ hybridization. For mitochondrial activity staining, embryos were directly embedded in OCT and snap frozen in liquid nitrogen and sectioned at 10µm using a cryostat.

**Histological and immunohistochemical analysis**

Histological sample processing and immunostaining was performed as previously described (*1*). The primary antibodies used in this study were: HIF1α (NB100-479, Novus Biologicals; USA); cTnT (CT3, Developmental Studies Hybridoma Bank; USA); BrdU (347580, BD Biosciences; USA); Cy3-conjugated Smooth Muscle Actin (C6198, Sigma Aldrich; USA) and GLUT1 (Cat. No. 07-1401, Millipore, USA). FITC-WGA (W32466, Life Technologies; USA) was used to stain cell membrane. Nuclei were stained with DAPI (Millipore; USA). Mitochondrial activity staining was performed on cryo-preserved fresh tissue sections as described previously (*1*).

**Quantification of histological and immunostained sections**

HE staining was quantified as described previously (*1*) using ImageJ (*2*). For fluorescence intensity analysis in cardiomyocyte nuclei, our own pipeline for CellProfiler software was employed (*3*). Briefly, cell nuclei were segmented and subsequently filtered by cTnT positive cytoplasmic staining. After filtration, HIF1α channel intensity was measured. For WGA-based area quantification, 25 cardiomyocytes in RV, LV and IVS were manually quantified. Only cardiomyocytes in cross-sectional area at the level of the nucleus were quantified.

**RNA extraction, cDNA synthesis and RT-qPCR**

RNA extraction from embryonic hearts, cDNA synthesis and quantitative PCR were performed as previously described (*1*). Primers are available under request.

**Probe synthesis and in situ hybridization**

General probe synthesis, purification and in situ hybridization steps were followed according with our previous protocol (*1*). For the probe synthesis, the following primers were used: *Glut1* 5’ GGACTTTGATGGCTCCAGAA 3’ and 5’ GAGTGTCCGTGTCTTCAGCA 3’, *Pdk1* 5’ CTGGGTTTGGTTACGGATTG 3’ and 5’ GCCAGCTACTCCACGTTCTT 3’ and *Ldha* 5’ GGAAGGAGGTTCACAAGCAG 3’ and 5’ CTGCAGTTGGCAGTGTGTCT 3’.

#### Electron microscopy and micrograph quantification

Embryonic hearts were processed for transmission electron microscopy following the standard procedures. Briefly, after overnight fixation in 3% glutaraldehyde/4% PFA, samples were refixed in 1% osmium tetroxide and embedded in epoxy resin. 60nm sections were counterstained with uranyl acetate and lead citrate and imaged using a JEOL JEM1010 (100 KV) transmission electron microscope. Control and mutant embryonic hearts from three independent litters were analyzed. For quantification of mitochondria and lipid droplets, ten images of compact myocardium and ten of trabeculae were taken at 5000x magnification. Mitochondria and droplets were counted manually by blinded observers using the ImageJ CellCounter plugin. Values were normalized to the total tissue area, in pixels, excluding extracellular areas in the image.

#### Protein extraction and Western Blot

Embryonic hearts were homogenyzed using RIPA buffer and a 25Ga needle in presence of protease and phosphatase inhibitors (Inhibitor cocktail (11697498001, Roche, Switzerland) and 1µM sodium ortovanadate). After clarification by centrifugation, protein concentration was measured using Pierce BA Protein Assay kit (23227, Thermo Scientific; USA) following manufacturer instructions. 30µg of protein were denaturized at 95ºC for 5 min, loaded on an 8% polyacrylamide SDS-PAGE gel and run at 120V for 90min. Subsequently, samples were transferred to a nitrocellulose membrane by wet transfer at 400mA for 2h. Membranes were blocked with 10% skimmed milk for 1h and incubated with primary antibodies O/N at 4ºC. Next day, membranes were washed in TBS-T buffer and incubated with the corresponding HRP-conjugated secondary antibodies (Dako, Denmark) at 1:5000 dilution for 1h at RT. After washing, signal was developed using ECL Primer Western Blotting Detection Reagent (Amersham; UK) and detected by a LAS-3000 imaging system (Fujifilm; USA). The primary antibodies used in this study were: anti-HIF1α (10006421, Cayman; USA) dilution 1:200, anti-tubulin (sc8035, Santa Cruz Biotech; USA) dilution1:1000 and anti-βActin (sc47778, Santa Cruz Biotech; USA) dilution 1:1000.

#### RNASeq and bioinformatics analysis of gene expression

RNASeq was performed as described previously (*1*). For analysis, only genes expressed at least at 1 count per million and at least in 2 samples were considered. Changes in gene expression were considered significant if associated with a Benjamini and Hochberg adjusted P value < 0.055. GO Term enrichment analysis was performed against Panther Biological Processes Database. Enriched GO terms were filtered by applying a P value threshold of 0.001 (default value) and manually grouped into ad hoc defined categories, as described in Table S2. The association between genes and enriched GO terms was represented graphically with GO plot (*4*).

#### Metabolomics untargeted profiling analysis by HPLC

Metabolites from single E12.5 embryonic hearts were extracted by polypropylene pestle homogenization in buffer containing 0.1% ammonium acetate:methanol (1:1 v/v) solution containing 5mM BHT. Proteins were precipitated by 1:1 methanol-ethanol. Blank sample was prepared following the same protocol only using the solvents. Quality Control (QC) samples were prepared by pooling 15µL of each sample homogenate. Supernatant and pellets were dried in speedvac speedvac (Savant SPD131DDA concentrator, Savant RVT5105 refrigerated vapor trap and OFP400 vacuum pump, ThermoFisher; USA) for 2h at RT. Pellets were stored at -80ºC for further proteomics analysis Metabolomics untargeted analysis was performed using a Ultimate 3000 HPLC system consisting of a degasser, two binary pumps, and thermostated autosampler, maintained at 8°C (ThermoFishe; USA) coupled to a LTQ Orbitrap XL™ Hybrid Ion Trap-Orbitrap Mass Spectrometer (ThermoFisher; USA). 5µL of the samples were injected onto a Merck SeQuant ZIC-HILIC column (150 × 1 mm, 3.5µ), which was thermostated at 45°C, and metabolites were eluting at 180µL/min with solvent A composed of water with 0.1% formic acid, and solvent B composed of acetonitrile with 0.1% formic acid. The gradient started from 90% to 25% of B in 15 min, keeping constant for 3 min and returned to starting conditions in 0.1 min, finally by keeping the re-equilibration at 90% of B for 11.9 min. Data were collected in positive and negative ESI ion modes in separated runs. The MS was operating in full scan mode from 70 to 1000 m/z at 60000 resolution. MS/MS spectra were collected in data-dependent mode *via* collision induced dissociation (CID) in the ion trap. Samples were analysed in a randomized order in two runs (first for positive and second for negative ion mode). QCs were analysed at the beginning, at the end and every six samples. Generated data were aligned using Compound Discoverer (ThermoFisher; USA) and signals were extracted and grouped into features y Metaboprofiler node. After filtering, keeping entities present in, at least, 75% of the samples, and showing CV% lower than 30% of QC value, were analyzed by the KNN method in MetaboAnalyst (http://www.metaboanalyst.ca/) and normalized by total protein content in each sample. Putative identification was performed using Ceu Mass Mediator (http://ceumass.eps.uspceu.es/).

**Proteomics analysis**

Proteins from pellets after metabolite extraction were pooled in groups of four, treated with 50mM iodoacetamide (IAM) and digested with trypsin using the Filter Aided Sample Preparation (FASP) digestion kit (Expedeon) (*5*) according to manufacturer's instructions. Dried peptides were labeled with iTRAQ-8plex according to manufacturer's instructions, desalted on OASIS HLB extraction cartidges (Waters Corp.), separated into 4 fractions using the high pH reversed-phase peptide fractionation kit (Thermo) and dried-down before MS analysis on an Orbitrap Fusion Tribrid mass spectrometer (Thermo Fisher Scientific, Bremen, Germany) (*6*). Peptide identification, quantification and systems biology analysis was performed as in (*6*–*8*). Significant abundance changes of proteins or homogeneous categories of KO mice compared to controls were detected at 1% FDR.

**Adult mice echocardiography and analysis**

5 months-old mice were anesthetized using 1.5% isoflurane at a flow rate of 1L/min. Once anesthetic plane was reached, cardiac images were acquired using a MS400 probe, at 30MHz for 2D and M mode images and 24MHz for Color and Pulsed Doppler modes, using an ultrasound scanner VEVO2100 (Visualsonics, Canada).

### SUPPLEMENTAL FIGURES AND FIGURE LEGENDS

#### Figure S1

**Deletion controls of *Hif1a/Nkx2.5* mutants. A)** Agarose electrophoresis showing PCR products from heart (H) and tail (T) tissue of E12.5 mutant embryos (*Hif1a*$^{f/f}$/*Nkx2*$^{rCre/+}$, lanes 1and 2) and controls (*Hif1a*$^{f/f}$/*Nkx2.5*$^{+/+}$, lanes 3 and 4). Top gel: floxed (615 bp) allele of the *Hif1a* gene. Middle gel: wild-type (264 pb) and Cre (583 bp) alleles of the Nkx2.5 gene. Bottom gel: processed *Hif1a* allele after Cre-mediated recombination (400bp) and unprocessed allele (1213bp). **B)** RT-qPCR quantification of *Hif1a,* exon 2 (floxed) from *Hif1a*, *Hif2a*, *Hif1b¸ Vhl, Phd3 and Vegf* transcripts in E14.5 *Hif1a-* mutant hearts. Bars (mean±SEM, n=3) represent fold induction relative to baseline expression in littermate controls (red line). *pvalue<0.05; **0.01<pvalue<0.05; ***pvalue<0.005, Student's t test. **C)** HIF1α immunofluorescence at E12.5 in control and mutant embryos (Dapi staining shows nuclei in blue, Troponin T in green and HIF1α in red). Scale bars, 20µm. **D)** Representative analysis of cardiomyocyte HIF1α nuclear protein expression intensity, quantified by immunohistochemical staining of heart sections from an E12.5 control embryo (green curve) and a *Hif1a*-null littermate (red curve).

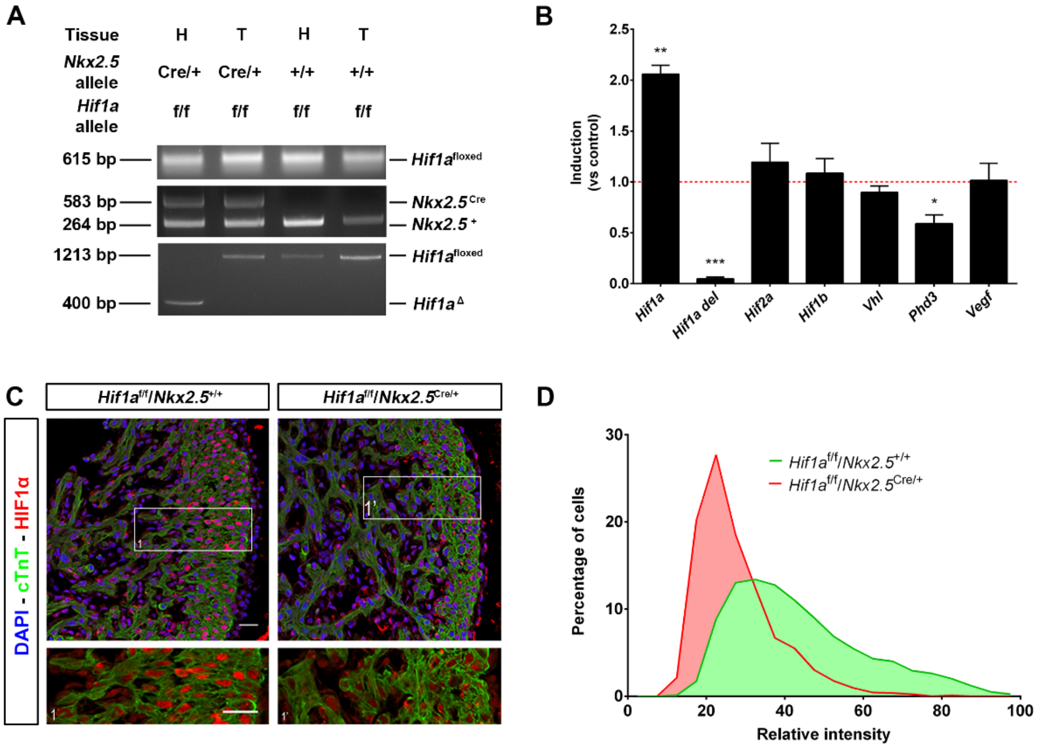

### Figure S2

**Embryonic analysis of *Hif1a/Nkx2.5* mutants at E14.5. A)** E14.5 control (*Hif1a*$^{f/f}$/*Nkx2*$^{r+/+}$, up) and mutant (*Hif1a*$^{f/f}$/*Nkx2.5*$^{Cre/+}$, down) embryos stained with hematoxylin and eosin (HE). Scale bars represent 100µm (overview) and 20µm (insets). **B)** HE quantification of ventricular walls and inter-ventricular septum width in E14.5 control (black bars) and mutant (white bars) embryos. **C)** LV magnifications of E14.5 control (up and black bar) and *Hif1a*-deficient (down and white bar) heart sections stained with wheat germ agglutinin (WGA) and quantification of cardiomyocyte cross-sectional area. **D)** Quantification of BrdU immunostaining, represented as percentage of BrdU$^{+}$ cells in the compact myocardium and trabeculae of E14.5 control (black) and *Hif1a/Nkx2.5* mutant (white) embryos. In all graphs, bars represent mean±SEM (n=3), Student's t test.

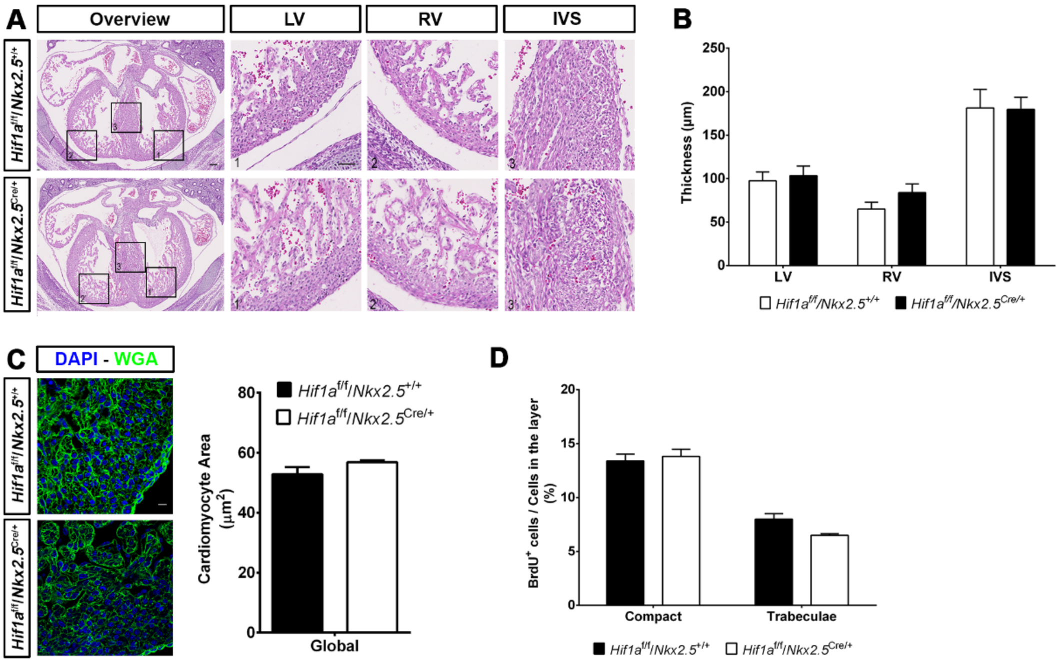

**Figure S3**

**Analysis of *Hif1a/cTnT* embryos at E14.5. A)** IVS magnifications of E14.5 cardiac sections in control (*Hif1a*$^{f/f}$/*cTnT*$^{+/+}$) and *Hif1a/cTnT* mutant (*Hif1a*$^{f/f}$/*cTnT*$^{Cre/+}$) embryos stained for HIF1α immunofluorescence (Dapi staining shows nuclei in blue, Troponin T in red and HIF1α in green). Scale bars, 20µm. **B)** E14.5 control and *Hif1a/cTnT* mutant embryos stained with hematoxylin and eosin (HE). Scale bars represent 100µm (overview) and 20µm (insets). **C)** RV magnifications of E14 control and *Hif1a/cTnT* embryos analyzed by in situ hybridization against *Glut1* (left), *Pdk1* (middle) and *Ldha* (right) mRNA expression. Scale bar represent 20µm. **D)** RT-qPCR analysis of glycolytic genes in E14.5 *Hif1a/cTnT* mutant ventricles. Bars (mean±SEM, n=3) represent fold induction relative to baseline expression in littermate controls (red line). Student's t test, *pvalue<0.05; ***pvalue<0.005. RV: right ventricle; LV: left ventricle; IVS: interventricular septum.

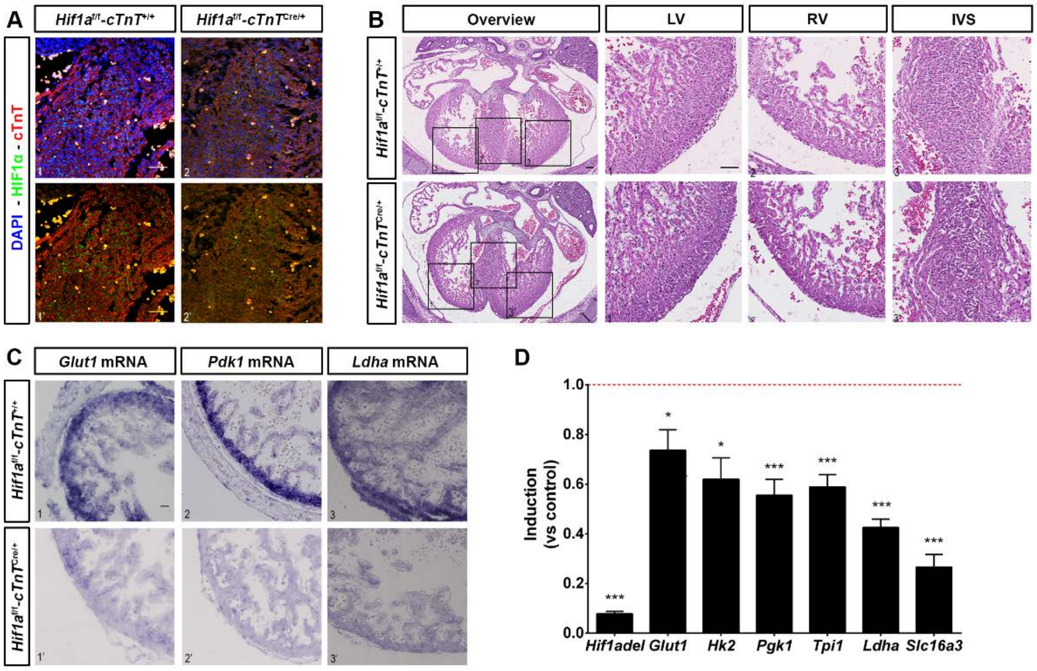

**Figure S4**

**Schematic representation of amino acid contributions to Krebs and Urea Cycles.** Schematic overview of the transcriptomics (mRNA expression, left column) and proteomics data (standardized protein quantifications, right column) of the re-wired metabolic pathways in the heart of *Hif1a/Nkx2.5* mutants (*Hif1a*$^{f/f}$*/Nkx2.5*$^{Cre/+}$) over control embryos (*Hif1a*$^{f/f}$*/Nkx2*$^{r+/+}$) at E12.5. Data are represented as individual heat maps for the transcript/protein of each pathway calculated as logarithmic Fold Change (logFC) and coded by color intensity following the scale at the bottom. ND indicates no detection. All 14406 expressed genes and 4276 quantified proteins were considered for this analysis.

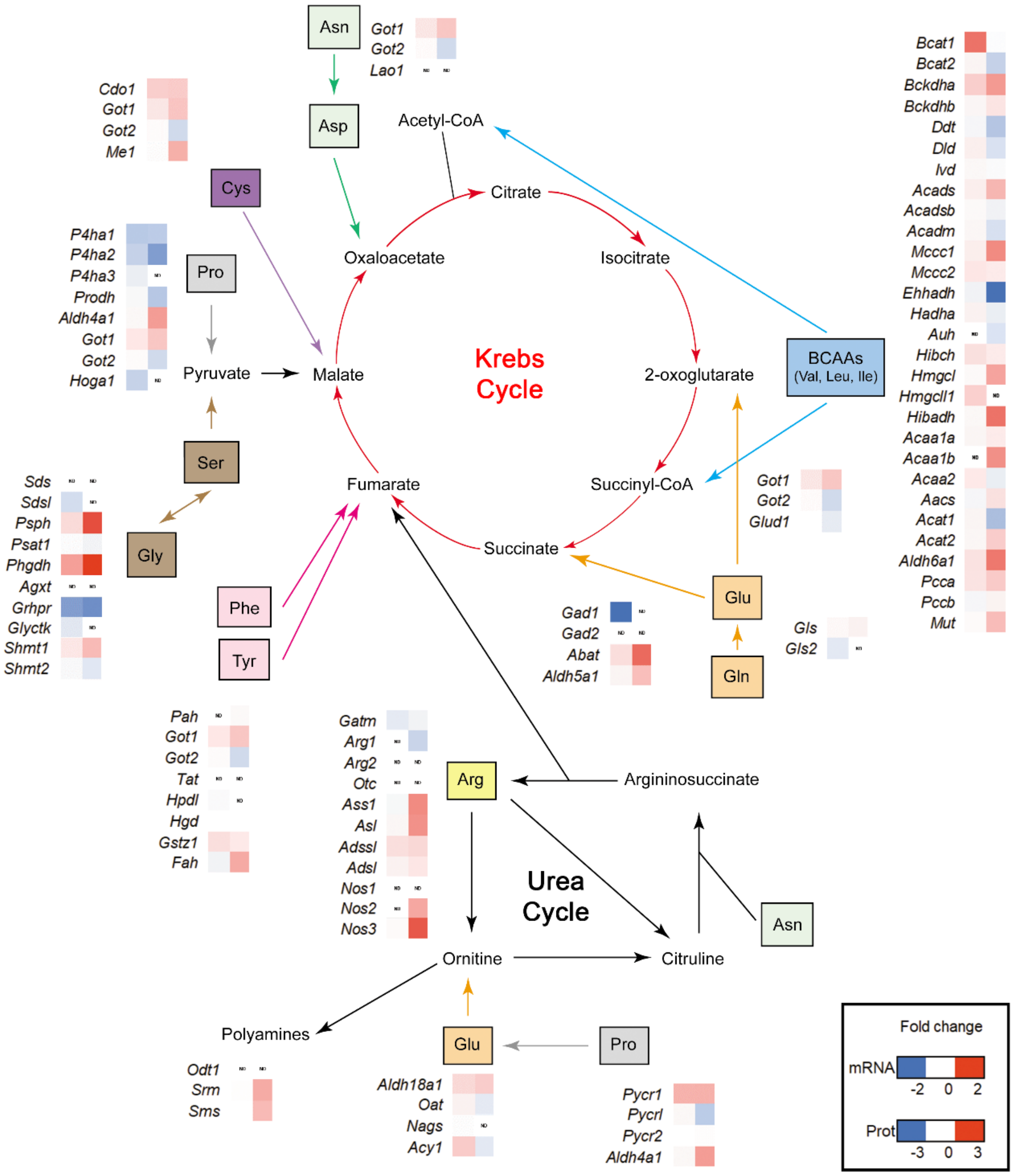

**FigureS5**

**Morphological and functional echocardiographic parameters in *Hif1a/Nkx2.5* adult mutants.** Quantification of IVS thickness **(A)**, LV posterior wall thickness **(B)**, LV volume **(C)** and diastolic function E/A **(D)** in control (*Hif1a*$^{f/f}$/*Nkx2*$^{r+/+}$, black bars n=9) and *Hif1a/Nkx2.5* (*Hif1a*$^{f/f}$/*Nkx2.5*$^{Cre/+}$, white bars n=11) at 5 months of age. Bars represent mean±SEM, Student's t test. IVS: interventricular septum; LV: left ventricle; E/A: E wave/A wave.

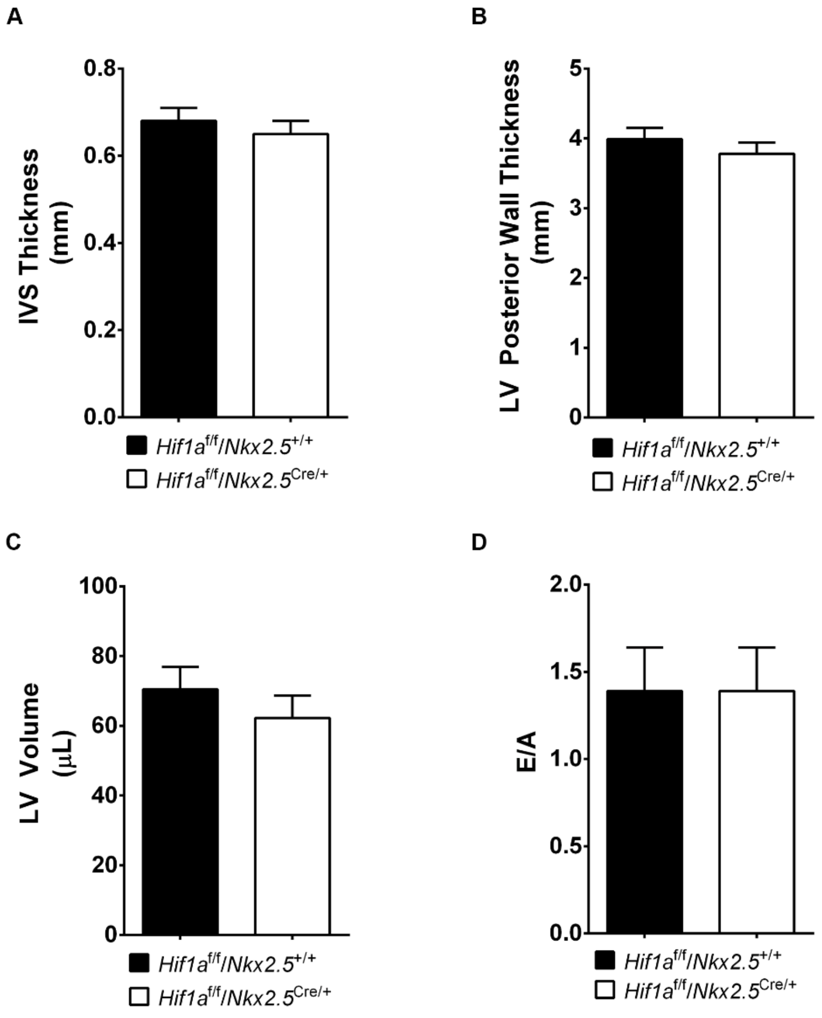

### SUPPLEMENTAL TABLES AND SUPPORTING INFORMATION

**Table S1**
201 genes detected as differentially expressed (DEG) in the Hif1a$^{flox/flox}$/Nkx2.5$^{Cre/+}$ versus Hif1a$^{flox/flox}$/Nkx2.5$^{+/+}$ contrast, with Benjamini-Hochberg adjusted p_value < 0.055. There are 83 UP-regulated genes and 118 DOWN-regulated genes. For each gene, the table describes Ensembl gene ID, MGI symbol, average expression value in Hif1a$^{flox/flox}$/Nkx2.5$^{Cre/+}$ and Hif1a$^{flox/flox}$/Nkx2.5$^{+/+}$ conditions, log fold of the expression change between the two conditions and its associated p-value, and gene description.

**Table S2**
Over-represented Biological Process GO terms associated to the collection of 201 differentially expressed genes (adjusted p_value < 0.055), as detected with GOrilla, with p_value < 0.001. The table describes, for each GO term, the number of mapped annotated genes in the complete set of expressed genes, the number of mapped annotated genes in the target set, its fold enrichment and p value.

**Table S3**
Over-represented Reactome pathways, Panther pathways and Biological Process GO terms associated to the collection of 201 differentially expressed genes (adjusted p_value < 0.055), as detected with PANTHER, with p_value < 0.05. The table describes, for each pathway or GO term, the number of mapped annotated genes in the complete set of expressed genes, the number of mapped annotated genes in the target set, its fold enrichment and p value.
